# Elucidation of the anti-inflammatory mechanism of isoliquiritigenin from *Glycyrrhiza uralensis* using activity-based protein profiling

**DOI:** 10.64898/2026.05.05.722967

**Authors:** Hina Sakai, Mi Hwa Chung, Taiki Nakaya, Katsuya Ohbuchi, Yosuke Isobe, Makoto Arita, Kazuya Tsumagari, Koshi Imami, Takatsugu Hirokawa, Hiroshi Tsugawa

**Affiliations:** Department of Biotechnology and Life Science, Tokyo University of Agriculture and Technology, 2-24-16 Naka-cho, Koganei, Tokyo, 184-8588 Japan; Tsumura Kampo Laboratories, Tsumura & Co, Ami, Ibaraki 300-1192, Japan; Division of Physiological Chemistry and Metabolism, Graduate School of Pharmaceutical Sciences, Keio University, 1-5-30 Shibakoen, Minato-ku, Tokyo, 105-8512 Japan; RIKEN Center for Integrative Medical Sciences, 1-7-22 Suehiro-cho, Tsurumi-ku, Yokohama, Kanagawa, 230-0045, Japan; Division of Biomedical Science, Faculty of Medicine, University of Tsukuba, 1-1-1 Tennodai, Tsukuba, Ibaraki, 305-8577 Japan; RIKEN Center for Sustainable Resource Science, 1-7-22 Suehiro-cho, Tsurumi-ku, Yokohama, Kanagawa 230-0045, Japan; Graduate School of Pharmaceutical Sciences, Kyoto University, 46-29 Yoshida-Shimoadachi-cho, Sakyo-ku, Kyoto 606-8501, Japan

**Author notes:** **Corresponding Author** Hiroshi Tsugawa.

## Abstract

*Glycyrrhiza uralensis* is a widely used medicinal plant present in more than 70% of Kampo formulations in Japan owing to its diverse pharmacological activities, including immunomodulatory, antitumor, and antioxidant effects. Isoliquiritigenin (ILG), a major chalcone constituent of *G. uralensis*, exhibits anti-inflammatory activity; however, its molecular mechanism remains unclear. Here, we employed an activity-based protein profiling approach to identify the molecular targets of ILG. Given that the α,β-unsaturated carbonyl moiety of ILG can covalently react with reactive cysteine residues via nucleophilic addition, we used an iodoacetamide-based probe to globally profile cysteine-reactive proteomes. The comparative analysis between ILG- and vehicle-treated RAW 264.7 macrophages identified cysteine 65 (Cys65) of lipocalin-type prostaglandin D_2_ synthase (L-PGDS) as a potential covalent target. ILG treatment did not alter L-PGDS expression levels, indicating that reduced probe labeling reflects direct covalent competition rather than changes in expression. Consistently, ILG significantly suppressed prostaglandin D_2_ (PGD_2_) production, comparable to the selective L-PGDS inhibitor AT-56. Although both ILG and AT-56 reduced interleukin-6 expression, ILG exerted a stronger inhibitory effect. Our results demonstrate that covalent inhibition of L-PGDS and subsequent suppression of PGD_2_ production represent a key mechanism underlying the anti-inflammatory activity of ILG.

## Introduction

Traditional herbal medicine, known as Kampo, was introduced to Japan from China during the fifth and sixth centuries and has since evolved uniquely, adapting to the local climate, soil conditions, and human constitution.^1^ By 2025, 148 Kampo formulations had been approved as prescription drugs and are widely used in clinical practice for the treatment of inflammatory, metabolic, neurological, and gastrointestinal disorders. These formulations typically consist of multiple herbal components and are characterized by a “multicomponent, multitarget” mode of action. Although numerous active compounds have been identified, their pleiotropic biological effects and complex phytochemical interactions complicate efforts to attribute *in vivo* efficacy to individual molecules. Among Kampo medicinal plants, species of the genus *Glycyrrhiza* (licorice) play a central role. Licorice is included in approximately 70% of Kampo prescriptions and has been used in East Asian medicine for centuries. Commercial preparations are primarily derived from the dried roots and stolons of *Glycyrrhiza uralensis* or *Glycyrrhiza glabra*. More than 300 phytochemicals have been characterized in licorice,^2^ including triterpenoid saponins, flavonoids, chalcones, and coumarins, many of which exhibit antioxidant, anti-inflammatory, antiviral, and anticancer activities.^3-5^ This chemical diversity likely underlies its long-standing use as a “harmonizing” component in traditional formulations.

Several licorice-derived compounds modulate innate immune responses. Glycyrrhizin (GL) and isoliquiritigenin (ILG), for example, attenuate pro-inflammatory signaling in macrophages, including the suppression of interleukin (IL)-6 production under lipopolysaccharide (LPS) stimulation.^6^ Although both compounds modulate Toll-like receptor 4 (TLR4) signaling, their mechanisms of action differ. While GL inhibits TLR4 dimerization and LPS binding to the TLR4/MD-2 complex, ILG modulates TLR4 signaling without directly blocking LPS binding.^6,7^ Moreover, a multi-omics approach has revealed that ILG induces distinct metabolic and phosphoproteomic profiles compared to crude licorice extract, suggesting that ILG contributes unique bioactivities.^8^

ILG is a representative chalcone-type flavonoid extensively studied for its anti-inflammatory properties, including the inhibition of nitric oxide (NO), cytokine, and prostaglandin production.^9,10^ These activities depend on the presence of the α,β-unsaturated carbonyl moiety characteristic of the chalcone scaffold, as reduction of this electrophilic double bond diminishes biological activity. The electrophilic nature of this moiety enables covalent modification of nucleophilic cysteine residues via Michael addition.^11, 12^ Based on this chemical reactivity, we hypothesized that ILG exerts its anti-inflammatory effects, at least in part, through covalent modification of functionally important cysteine residues in immune-related proteins. To test this hypothesis, we employed activity-based protein profiling (ABPP), a chemoproteomic strategy that enables global identification of proteins containing reactive cysteine residues.^13^ In combination with tandem mass tag (TMT)-based quantitative proteomics, ABPP allows systematic evaluation of covalent target engagement at the proteome scale.^14^ Here, we applied a quantitative ABPP workflow to identify cysteine-reactive protein targets of ILG in macrophages. Candidate targets were further validated using *in silico* interaction analysis and complementary biochemical assays. This integrated approach provides mechanistic insight into the pharmacological actions of ILG and establishes a general framework for elucidating the molecular basis of bioactive phytochemicals in traditional herbal medicine.

## Results

### Proteome-wide screening for covalent binding targets of ILG

Quantitative tandem mass tag activity-based protein profiling (TMT-ABPP) was performed to identify ILG-binding proteins harboring reactive cysteine residues in macrophages. ILG contains a chalcone scaffold with an α,β-unsaturated carbonyl moiety capable of undergoing Michael addition with nucleophilic cysteine residues, supporting its potential to form covalent adducts with protein targets (**Figure 1A and 1B**).^9,10,12^ Based on this electrophilic property, RAW264.7 macrophages were treated with ILG (10 µM or 30 µM) or vehicle (DMSO), followed by labeling with the cysteine-reactive probe desthiobiotin-iodoacetamide (DB-IA). After trypsin/LysC digestion, DB-IA-conjugated peptides were enriched using streptavidin beads and analyzed by a nanoLC–MS/MS system (**Figure 1C**). In total, 2,688 unique cysteine-containing peptides were quantified (**Figure 1D and 1E**). Competition was assessed using the log_2_-transformed peptide intensity ratio (DMSO vs ILG), where decreased DB-IA labeling in ILG-treated samples indicates covalent engagement. Among the identified targets, lipocalin-type prostaglandin D2 synthase (L-PGDS) showed the strongest competition in both the 10 µM and 30 µM ILG treatment conditions, indicating pronounced suppression of DB-IA labeling in the presence of ILG. L-PGDS catalyzes the conversion of prostaglandin H2 (PGH_2_) to prostaglandin D2 (PGD_2_) within the arachidonic acid cascade. The MS^2^ spectrum analysis localized DB-IA modification to 65^th^ cysteine (Cys65) in L-PGDS, and the MS^3^-based TMT quantification confirmed dose-dependent competition at this site (**Figure 1F, 1G, and S1**). Cys65 is a known catalytic residue essential for L-PGDS enzymatic activity, suggesting that ILG directly targets the functional active site. We next examined proteins exhibiting competition ratios greater than two. Several enriched targets are functionally linked to prostaglandin signaling pathways. These include the aldo-keto reductase family 1 member B1 (AKR1B1), which reduces PGH_2_- or PGD_2_-derived intermediates,^15^ and the adaptor protein phosphotyrosine interacting with the PH domain and leucine zipper 2 (APPL2), an adaptor protein that mediates phosphoinositide 3-kinase (PI3K)–Akt signaling downstream of prostaglandin D receptors (DP1/DP2).^16^ In addition, RhoC, a Rho-family GTPase involved in PI3K/Akt regulation,^17^ was also enriched. These proteins are associated with metabolic and signaling processes downstream of PGD_2_, suggesting that ILG perturbs the L-PGDS-centered prostaglandin network in a coordinated manner. In addition, oxysterol-binding protein 8 (ORP8), which has been implicated in inflammatory signaling and apoptosis,^18^ was identified, indicating that ILG may modulate additional inflammation-related pathways.

**Figure 1.**
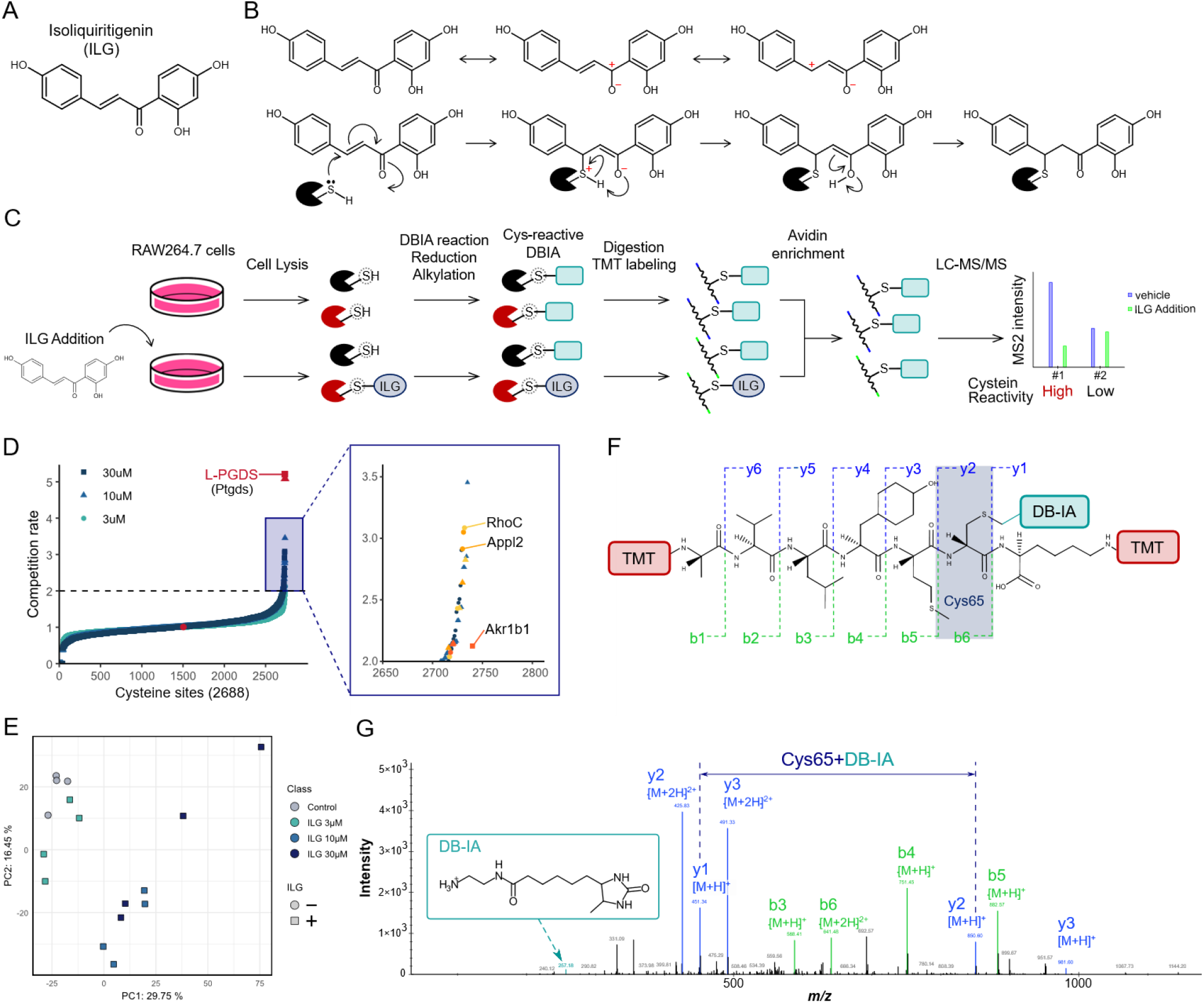
Overview of tandem mass tag–based activity-based protein profiling (TMT-ABPP) used in this study. (A) Chemical structure of isoliquiritigenin (ILG). (B) Schematic illustration of the covalent reaction between ILG and cysteine residues via Michael addition. Workflow of TMT-ABPP–based chemoproteomics analysis. (D) Distribution of competition rates across 2,688 quantified cysteine-containing peptides. Competition rate is defined as the log_2_ intensity ratio (DMSO vs ILG). (E) Principal component analysis (PCA) score plot based on enriched peptide abundances. (F) Structure of the L-PGDS peptide modified with TMT tags and desthiobiotin-iodoacetamide (DB-IA), highlighting the Cys65 residue. (G) Representative MS2 spectrum of the L-PGDS peptide shown in (F), with reporter ion–based quantification obtained from MS3 analysis.

PGD_2_ is a lipid mediator with context-dependent pro- and anti-inflammatory roles.^19^ It is synthesized from arachidonic acid via cyclooxygenase (COX)-mediated conversion to PGH_2_, followed by isomerization by L-PGDS (**Figure 2A**). Two PGD synthase isoforms exist: lipocalin-type (L-PGDS) and hematopoietic-type (H-PGDS), which differ in sequence and tissue distribution.^20^ Although L-PGDS is expressed at low basal levels in resting macrophages, its expression increases markedly under LPS-induced inflammatory conditions, accounting for approximately two-thirds of total PGD_2_ production in RAW264.7 cells.^21^ Structurally, L-PGDS contains two binding pockets. Pocket 1 includes Cys65 within the catalytic site responsible for PGD_2_ synthesis,^22, 23^ whereas Pocket 2 forms a hydrophobic cavity composed of aromatic residues (Phe34, Phe39, Trp43, Tyr105, and Tyr149) that accommodates ligands such as retinoic acid. Notably, L-PGDS possesses an unusually large internal cavity among lipocalins, enabling multiple ligand-binding modes. This structural feature likely underlies its broad ligand selectivity and functional versatility compared with other lipocalin family members.^24^

**Figure 2.**
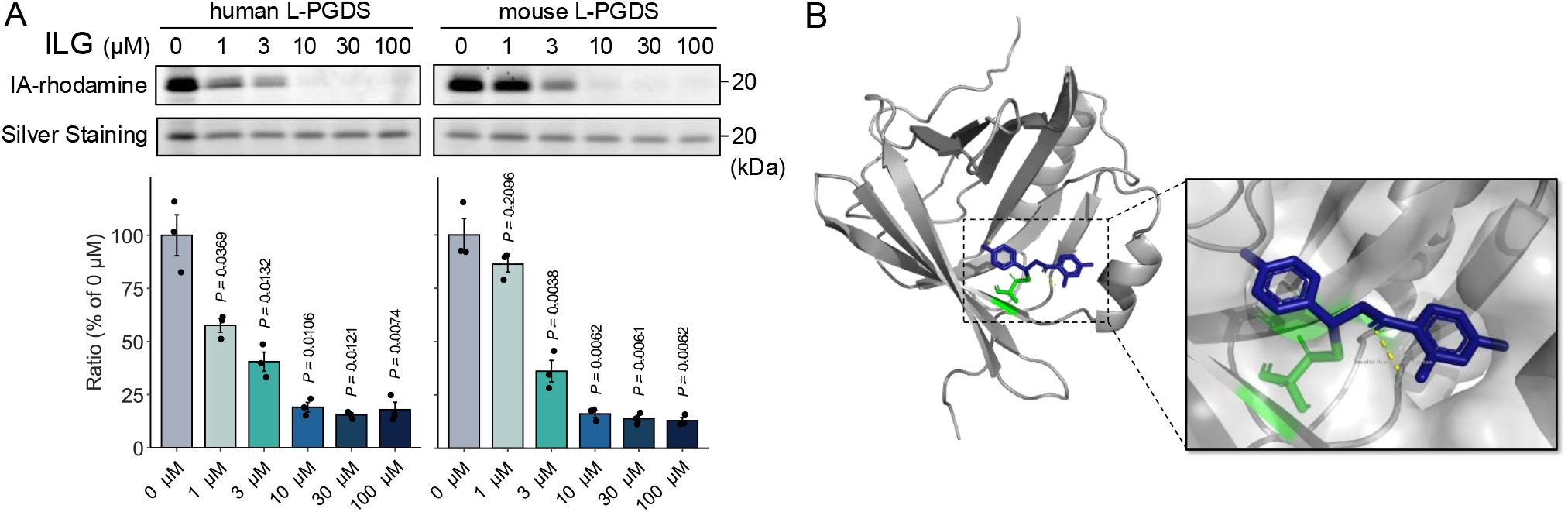
Covalent interaction between ILG and L-PGDS at Cys65. (A) Gel-based ABPP validation of ILG binding to L-PGDS. Recombinant human and mouse L-PGDS proteins were preincubated with ILG (0, 1, 3, 10, 30, and 100 µM) prior to labeling with IA-rhodamine. Proteins were separated by SDS–PAGE, and probe labeling was visualized by in-gel fluorescence. (B) Covalent docking model of ILG bound to L-PGDS (PDB ID: 2E4J) at Cys65. L-PGDS is shown in gray, Cys65 in green, and ILG in blue, represented as cartoon and stick models.

### Covalent bond formation between ILG and L-PGDS Cys65

Based on the ABPP results, we performed gel-based ABPP using a rhodamine-functionalized iodoacetamide probe (IA-rhodamine) to validate whether L-PGDS is a direct covalent target of ILG. Recombinant human and mouse L-PGDS proteins were synthesized using a cell-free expression system. ILG dose-dependently reduced IA-rhodamine labeling of both human and mouse L-PGDS (Figure 2B), indicating direct competition for cysteine modification. Notably, human L-PGDS exhibited a pronounced reduction in labeling even at low ILG concentrations (1 µM). In contrast, mouse L-PGDS showed a stronger effect at higher concentrations (10 and 30 µM), consistent with the dose-dependent trends observed in the proteome-wide ABPP analysis.

Furthermore, we performed covalent docking, molecular dynamics (MD) simulations, and binding free energy calculations using the CovDock module implemented in the Schrödinger Suite to evaluate the binding pose and covalent binding potential of ILG toward L-PGDS. Given the availability of multiple L-PGDS conformations in the Protein Data Bank, 15 structures were subjected to pairwise root-mean-square deviation (RMSD) based clustering, and a representative structure with a low average RMSD was selected as the initial model. Initial CovDock simulations, specifying a Michael addition between the electrophilic moiety of ILG and Cys65, failed to generate a pre-reactive pose compatible with the canonical reaction geometry. Specifically, the thiol proton of Cys65 was not appropriately oriented for abstraction, which is required for nucleophilic attack. To overcome this limitation, we performed non-covalent docking using YASARA program, followed by short MD simulations to sample a broader conformational space within the L-PGDS binding pocket. From these simulations, candidate poses were selected based on geometric criteria consistent with nucleophilic attack and proton abstraction. These refined poses were then used as input for a second round of CovDock simulations. During the docking procedure, the cysteine-reactive pocket was constrained to preserve the binding-site geometry, and covalent reaction parameters were defined with Cys65 as the reactive residue and Michael addition as the reaction type. Binding energies were evaluated using MM-GBSA, which accounts for solvation effects and provides a reliable approximation for ligand–protein interactions in large systems. This integrated computational workflow identified energetically favorable conformations in which ILG is optimally positioned toward Cys65, enabling access to the electrophilic center required for proton abstraction and covalent bond formation (**Figure 2C**).

### ILG suppresses PGD_2_ production and inflammatory responses via L-PGDS targeting

To determine whether ILG affects L-PGDS protein abundance, we performed western blot analysis in the presence and absence of ILG (**Figure 3A**). No significant changes in L-PGDS expression were observed, indicating that the reduced probe labeling in ABPP experiments reflects direct competitive binding rather than altered protein levels. L-PGDS catalyzes a key step in the arachidonic acid cascade, generating the lipid mediator PGD_2_. We therefore examined whether ILG-mediated targeting of L-PGDS affects PGD_2_ production. PGD_2_ levels were quantified by LC–MS in LPS-stimulated RAW264.7 cells treated with DMSO, ILG, or AT-56, a selective L-PGDS inhibitor that covalently targets Cys65 (Figure 3B).^25^ LPS stimulation increased PGD_2_ production, which was significantly suppressed by both ILG and AT-56. These results support that ILG inhibits L-PGDS enzymatic activity through competitive targeting of Cys65.

**Figure 3.**
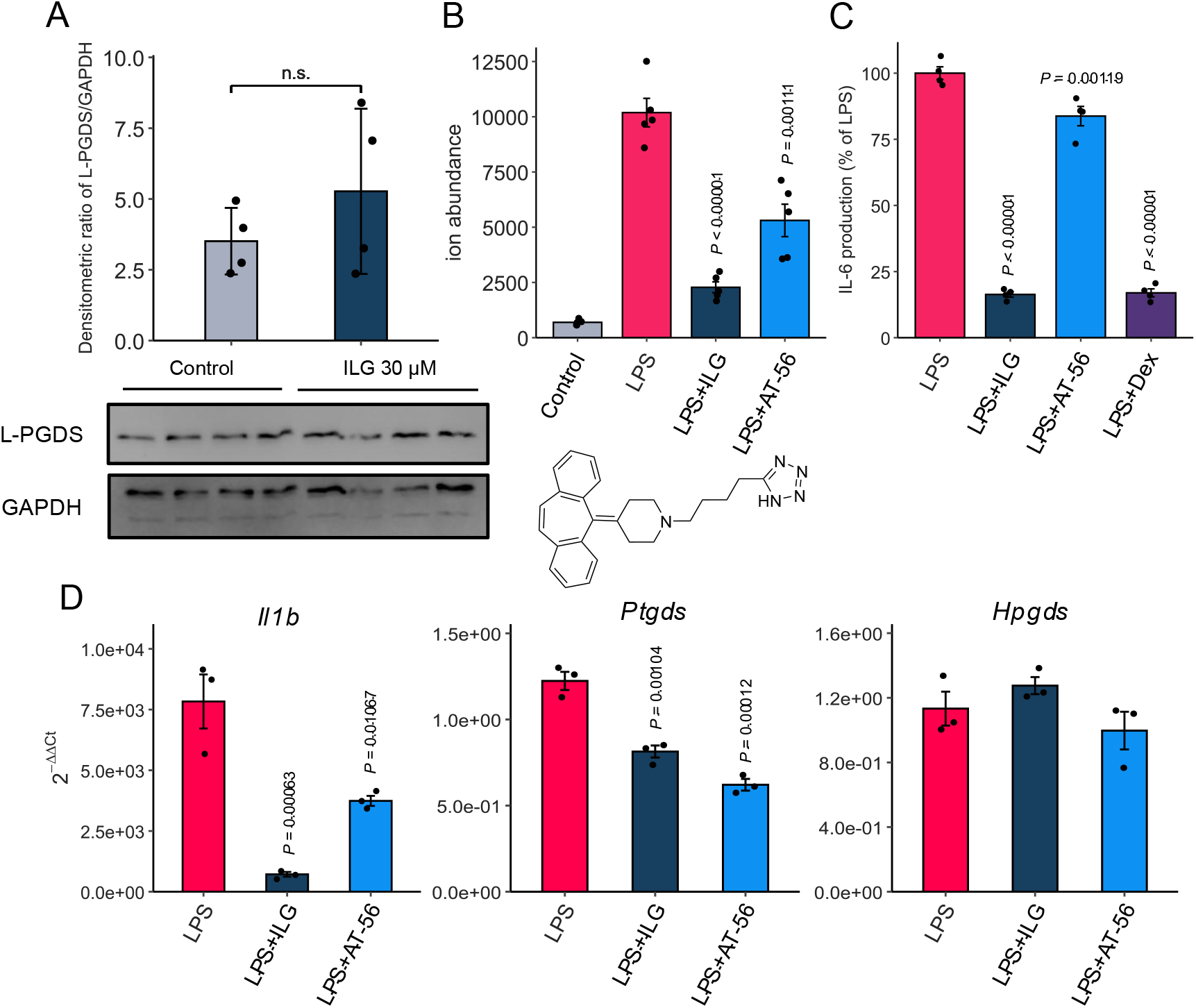
Functional effects of ILG on L-PGDS activity and inflammatory responses in RAW264.7 cells. (A) L-PGDS protein expression levels assessed by western blotting and normalized to GAPDH. (B) Ion abundance of PGD_2_ in RAW264.7 cells with or without preincubation with ILG (30 µM) or AT-56 (100 µM). PGD_2_ peak height was quantified by LC– MS and normalized to an internal standard. The chemical structure of AT-56 is shown. The values are presented as means ± standard error of the mean (SEM) with n = 5 biologically independent samples. (C) IL-6 production in RAW264.7 cells treated with ILG, AT-56, or dexamethasone (DEX, 1 µM) following 24 h stimulation with LPS (100 ng/mL). IL-6 levels were quantified by ELISA. Data are presented as mean ± SEM (n = 4 biological replicates). mRNA expression levels of IL-1β, L-PGDS, and H-PGDS in RAW264.7 cells treated with ILG or AT-56 following 3 h of stimulation with 100 ng/mL LPS. Statistical significance in (A– D) was evaluated by one-way ANOVA followed by Dunnett’s test, with comparisons made to the LPS-treated group.

PGD_2_ signaling exerts context-dependent immunological effects. For example, activation of the G protein–coupled receptor CRTH2 promotes Th2 cell migration and allergic inflammation, whereas signaling through DP receptors suppresses immune cell activation and migration.^26,27^ Thus, we evaluated IL-6 expression levels to examine the downstream biological implications of PGD_2_ reduction **(Figure 3C**). IL-6 levels were significantly decreased in both ILG- and AT-56-treated cells, with ILG showing a stronger inhibitory effect. Dexamethasone (DEX) was used as a positive control. In contrast to DEX, which broadly suppresses inflammatory gene transcription, ILG could appear to modulate inflammation through lipid mediator regulation. Consistent with these findings, IL-1β mRNA expression was induced by LPS and significantly reduced by ILG treatment (**Figure 3D**). AT-56 also reduced cytokine levels, although to a lesser extent. In addition, mRNA expression of L-PGDS and H-PGDS was examined (Figure 3D). Both genes were upregulated by LPS stimulation; however, ILG and AT-56 selectively reduced L-PGDS expression without affecting H-PGDS. These results demonstrate that ILG targets L-PGDS to suppress its enzymatic activity and PGD_2_ biosynthesis, thereby attenuating downstream inflammatory responses, including IL-6 and IL-1β production.

## Discussion

In this study, we applied activity-based protein profiling (ABPP) to ILG-treated RAW264.7 cells to identify covalent molecular targets and uncovered Cys65 of L-PGDS as a primary site of covalent engagement. Because L-PGDS protein abundance was not altered by ILG treatment, the reduced peptide signal observed in ABPP is most consistent with direct covalent modification leading to functional inhibition rather than transcriptional or proteolytic regulation. Furthermore, ILG suppressed PGD_2_ production under LPS-induced inflammatory conditions and reduced IL-6 expression, supporting a functional link between L-PGDS inhibition, lipid mediator regulation, and downstream cytokine responses. However, the role of L-PGDS and PGD_2_ signaling in inflammation is highly context-dependent. For instance, hematopoietic PGD synthase knockout mice exhibit exaggerated inflammation and delayed resolution in acute peritonitis models.^28^ Moreover, activation of DP1 receptors generally exerts anti-inflammatory effects via cAMP signaling, whereas CRTH2 (DP2) activation promotes eosinophil recruitment and Th2-mediated inflammation.^29^ In macrophages, PGD_2_ has also been reported to function as an autocrine regulator that limits excessive cytokine release during acute activation.^30^ In this context, the observed reduction in both PGD_2_ and IL-6 following ILG treatment may reflect suppression of lipid mediator–driven cytokine amplification. Notably, ILG exhibited approximately twofold stronger inhibition of PGD_2_ production and nearly fourfold greater suppression of IL-6 expression compared with AT-56. While AT-56 is a reversible inhibitor that binds near the active site,^25^ ILG covalently modifies Cys65, which may confer more sustained target engagement and explain its enhanced biological effects. These findings suggest that ILG acts as a covalent modulator of L-PGDS, distinct from classical reversible inhibitors, and that this mode of engagement translates into amplified anti-inflammatory outcomes.

Nevertheless, several limitations should be considered. First, the ABPP experiments were conducted in an immortalized macrophage cell line (RAW264.7), which does not fully recapitulate the complexity of primary or tissue-resident macrophages in vivo. Validation in primary murine macrophages, human monocyte-derived macrophages, and *in vivo* inflammation models (e.g., LPS-induced peritonitis) would strengthen the physiological relevance of these findings.^31, 32^ Second, ABPP primarily captures covalent interactions and may overlook transient or non-covalent binding events, suggesting that ILG-mediated cytokine suppression may not be solely explained by L-PGDS inhibition. Rescue experiments using exogenous PGD_2_ or receptor-specific modulators (DP1 vs DP2/CRTH2) will be important to dissect pathway specificity. In addition, the downstream metabolite 15-deoxy-Δ^12^,^14^-PGJ_2_, a ligand of peroxisome proliferator-activated receptor γ (PPARγ), may be altered following L-PGDS inhibition, which could indirectly influence cytokine transcription.^28,30^ Therefore, comprehensive metabolomic profiling would be crucial to determine whether the observed suppression of IL-6 results from a broader shift in eicosanoid signaling.

From a medicinal chemistry perspective, the covalent interaction between ILG and L-PGDS provides a framework for developing electrophilic modulators of prostaglandin metabolism. Given the tight coupling between prostaglandin and cytokine signaling in macrophages, it will be important to assess whether ILG selectively targets the PGD_2_ pathway or more broadly affects arachidonic acid metabolism, including PGE_2_ (mPGES-1) and PGI_2_ (PGIS) pathways. In conclusion, we identify L-PGDS as a previously unrecognized molecular target of ILG and demonstrate that covalent modification at Cys65 suppresses PGD_2_ production and downstream inflammatory responses. This mechanism provides a new perspective on the anti-inflammatory activity of ILG by linking electrophilic small-molecule reactivity to lipid mediator regulation and highlights L-PGDS as a potential therapeutic target in macrophage-driven inflammatory diseases.

## Methods

### Cell culture

RAW264.7 cells with passage numbers below 20 were cultured in Dulbecco’s Modified Eagle Medium (DMEM; High Glucose) with L-glutamine and phenol red (Fujifilm Wako) supplemented with 10% fetal bovine serum (FBS) (EquaFETAL; Atlas Biologicals) and 1% penicillin–streptomycin solution (Fujifilm Wako). RAW264.7, derived from BALB/c mice and transformed with the Abelson murine leukemia virus, is a well-established murine macrophage-like model. Upon LPS-mediated stimulation, these cells reliably produce pro-inflammatory mediators, including NO and cytokines, such as tumor necrosis factor-α, IL-1β, and IL-6. This response is mediated largely through NF-κB and MAPK-mediated signaling pathways, making this cell line a widely accepted *in vitro* platform for screening anti-inflammatory activity. In this study, this cell line was used to evaluate the anti-inflammatory effects of ILG. Cells were grown under a humidified atmosphere with 5% CO_2_ at 37 °C in a 10 cm tissue culture dish (BMBio). The cells were split once a week by washing with phosphate-buffered saline (D-PBS(-); Fujifilm Wako), followed by treatment with trypsin-EDTA (Promega) for cell detachment. After detachment, trypsin was quenched by adding the growth medium. A total of 5.0 × 10^5^ cells/dish were grown in fresh growth medium in a new dish.

### Cell pretreatment for TMT-ABPP

Experiments were performed with 4 - 5.0 × 10^6^ cells plated in fresh growth medium in a 10 cm tissue culture dish 1 day before ABPP pretreatment and grown under a humidified atmosphere with 5% CO_2_ at 37 °C. Dimethyl sulfoxide (DMSO; Fujifilm Wako) or ILG (Tokyo Chemical Industry Co., Ltd.) was added to the growth medium (final concentrations of 3, 10, and 30 µM), and the mixture was incubated under 5% CO_2_ at 37 °C for 1 h. The cells were washed with PBS and harvested using a cell scraper in 1 mL of PBS. Then, the cell suspension was spun down at 200 × *g* for 5 min at 4 °C, and the supernatant was aspirated. The cell pellet was snap-frozen in liquid nitrogen and stored at −80 °C for later use.

Cell pellets were lysed in PBS and sonicated three times using a probe sonicator at high amplitude for 3 s each. Samples were spun down for 5 min at 1,000 × *g* at 4 °C. The supernatant was collected, and proteins were analyzed using the Bicinchoninic Acid assay (BCA assay). Samples were diluted in PBS to 50 µg of protein per sample and incubated with 50 mM activity-based probe DB-IA in DMSO for 1 h at room temperature (RT). Excess DB-IA and disulfide bonds were quenched and reduced using 5 mM dithiothreitol (Fujifilm Wako, Osaka, Japan) for 30 min in the dark at RT. Subsequently, the reduced cysteine residues were alkylated with 20 mM iodoacetamide (Fujifilm Wako, Osaka, Japan) for 30 min in the dark at RT. Proteins were precipitated using MeOH/chloroform (CHCl_3_). A total of 400 µL of MeOH was added, followed by 100 µL of CHCl3 with thorough vortexing. After adding ultrapure water, the samples were centrifuged at 15,000 rpm for 3 min at RT, and the aqueous upper layer was removed. After adding MeOH (Fujifilm Wako), samples were centrifuged at 15,000 rpm for 3 min at RT, and protein pellets were air-dried. Protein pellets were resolubilized in 200 mM 4-(2-hydroxyethyl)-l-piperazine-propanesulfonic acid (EPPS; Thermo Fisher Scientific, USA) and digested with Trypsin/LysC (enzyme: protein = 1:25; Promega, USA) overnight at 37 °C using a ThermoMixer set to 1200 rpm. Tryptic peptides were incubated with 1 mg TMTpro 16plex reagent (Thermo Fisher Scientific) at RT for 1 h in the dark; the reaction was quenched by adding hydroxylamine at a final concentration of 5% (Sigma) for 15 min. All peptides were pooled and dried using SpeedVac. The mixed peptides were applied to streptavidin magnetic beads (Pierce, USA) overnight at 4 ºC, and were then eluted with 50% acetonitrile containing 0.1% TFA. The sample was further dried using SpeedVac, reconstituted with 1% formic acid, and desalted using SDB-RPS StageTips. Raw data were processed using MSFragger (v3.5), and source proteins for each peptide were identified.

### Analysis of TMT-labeled peptides using LC-MS/MS

A nano-LC-MS/MS system comprising an EASY-nLC 1200 pump (Thermo Fisher Scientific) and an Orbitrap Eclipse mass spectrometer (Thermo Fisher Scientific) equipped with a FAIMSpro interface (Thermo Fisher Scientific) was used. The mobile phases consisted of (A) 0.1% formic acid in water (A) and 0.1% formic acid in 80% acetonitrile. Peptides were loaded onto a self-made 20 cm fused-silica emitter (75 µm inner diameter, GL Sciences) packed with ReproSil-Pur C18-AQ (1.9 µm, Dr. Maisch, Ammerbuch, Germany) and separated using a 70 min linear gradient (5–10% B in 5 min, 10–40% B in 50 min, 40–99% B in 5 min, and 99% B for 10 min) at a flow rate of 300 nL/min. All MS1 spectra were acquired over the range of 400– 1500 *m/z* using the Orbitrap analyzer (resolution = 120,000, maximum injection time = “Auto,” automatic gain control = “Standard”). For the subsequent MS/MS analysis, precursor ions were selected and isolated in top-speed mode (cycle time = 1.5 s, isolation window = 0.7 *m/z*) with FAIMS compensation voltage (CV) stepping between two CVs, fragmented by collision-induced dissociation (CID; normalized collision energy = 35), and detected using the ion trap analyzer (turbo mode, maximum injection time = 35 ms, automatic gain control = “Standard”). The top ten most intense fragment ions were subjected to TMT reporter ion quantification using SPS-MS^3^ (HCD-normalized collision energy = 55).^33^ Samples were analyzed in quadruplicate with different FAIMS CV combinations (−30/–70, –40/–80, −50/–90, and –60/–100 V).

Raw data were processed using the FragPipe (version 19.0) suite with MSFragger (version 3.6), Philosopher (version 4.6.0), and IonQuant (version 1.8.9). Database searches were performed using the UniProt mouse proteome database (April 2022). Precursor and fragment ion mass tolerances were set to 20 ppm and 0.6 Da, respectively, with mass calibration and parameter optimization enabled. Strict trypsin cleavage specificity was applied, allowing up to two missed cleavages. Cysteine carbamidomethylation and TMTpro modification of the peptide N-terminus and lysine residues were set as fixed modifications, whereas oxidation of methionine and DB-IA modification (+239.16293 Da) of cysteine were allowed as variable modifications. The search results were filtered for a false discovery rate of <1% at the peptide spectrum match, ion, peptide, and protein levels.

### Recombinant protein synthesis

Recombinant Lipocalin-type prostaglandin d2 synthase (L-PGDS) and its mutant variants were synthesized using a wheat germ cell-free protein expression system (CellFree Sciences) in accordance with the manufacturer’s protocol. The full-length cDNA encoding L-PGDS was subcloned into the pEU vector incorporating an N-terminal 6×His tag. *In vitro* transcription was performed by combining the pEU construct with transcription buffer, NTP mix, RNase inhibitor, and SP6 RNA polymerase, followed by incubation at 37 °C for 6 h. The resulting mRNA was subjected to translation by supplementation with wheat germ extract WEPRO7240H, creatine kinase, and SUB-AMIX reagent, and the reaction mixture was incubated at RT for 20 h in the dark. The expressed recombinant proteins were subsequently purified by affinity chromatography using Nickel Sepharose resin (Cytiva) at 4 °C for 1 h, and eluted with buffer containing 500 mM imidazole (pH 7.5). Protein purity and expression were assessed by sodium dodecyl sulfate–polyacrylamide gel electrophoresis (SDS-PAGE).

### Gel-based ABPP

Recombinant purified protein (0.1 µg per sample) was pretreated with either DMSO (vehicle) or ILG at 37 °C for 30 min in 25 µL PBS and subsequently treated with 200 nM IA-rhodamine (Setareh Biotech) at RT for 1 h. The reaction was stopped by adding 4× reducing sodium dodecyl sulfate sample-loading buffer. After boiling at 95 °C for 5 min, the samples were separated using 10% SDS‒PAGE gels. Probe-labeled proteins were analyzed by in-gel fluorescence using a ChemiDoc MP (Bio-Rad).

### Western blotting

Cells were washed twice with D-PBS, and cellular proteins were solubilized and extracted using radioimmunoprecipitation assay (RIPA) buffer (Fujifilm Wako) on ice. The protein concentration was quantified using a bicinchoninic acid (BCA) assay, and equal amounts of protein (5 µg per lane) were loaded onto SDS‒PAGE gels and electrophoretically transferred to cellulose nitrate membranes. The membranes were blocked with 5% skim milk in TBS-T and incubated overnight at 4 °C with an anti–L-PGDS antibody (Abcam) as the primary antibody, followed by a 1 h incubation with a secondary antibody (Promega). Proteins were visualized using the chemiluminescent reagent ImmunoStar LD (Fujifilm Wako). Relative protein expression was analyzed using a Vilber FUSION Bio Imaging system (M&S Instruments Inc.) and normalized to glyceraldehyde-3-phosphate dehydrogenase (*Gapdh*) expression. The following antibodies were used for immunoblotting: anti–prostaglandin D synthase (lipocalin) antibody (ab182784; Abcam) and anti-GAPDH antibody (clone 14C10; Cell Signaling Technology).

### Lipid mediator analysis using LC-MS/MS

RAW264.7 cells were washed twice with D-PBS, resuspended in D-MEM, and seeded onto a 96-well plate (Corning) at 37 °C for 2 h and then treated with DMEM containing vehicle or ILG for 1 h. Thereafter, LPS was added to the culture medium. After incubation for 6 h at 37 °C, cells were washed a third time with prewarmed HBSS (Fujifilm Wako) and incubated with HBSS for 30 min. The medium was collected and centrifuged to remove cell debris. Lipids were extracted by adding methanol directly to the wells and incubating on ice. The medium and cell extracts were purified by solid-phase extraction using a MonoSpin C_18_-AX spin column (GL Sciences). The column was first equilibrated with MeOH and Milli-Q water, followed by sample application. After washing twice with Milli-Q water and 60% MeOH, prostaglandins were eluted with 90% MeOH and 2% acetic acid. The eluate was stored at −80 °C until analysis. LC-MS/MS-based analysis (LCMS-9030, Shimadzu) was performed using an Acquity UPLC BEH C_18_ column (1.0 mm × 150 mm, 1.7 µm particle size; Waters). Samples were separated using a linear gradient of Milli-Q water/acetic acid (1000:1, v/v) and acetonitrile/MeOH (4:1, v/v) for 14 min. MS/MS analysis was conducted in negative ion mode, and lipid metabolites were identified and quantified by multiple reaction monitoring. Calibration curves were prepared at five points ranging from 1 to 50 µM.

### Enzyme-linked immunosorbent assay (ELISA) for IL-6 production

IL-6 levels in RAW264.7 cells were measured using an ELISA kit (BD Biosciences), with four biological replicates per condition. Conjugate binding to the wells was analyzed by measuring absorbance using a Synergy™ HTX plate reader (BioTek). Dexamethasone (DEX), a potent synthetic glucocorticoid receptor agonist known to suppress NF-κB–mediated inflammatory gene expression, was used as the positive control.

### Molecular modeling and MD simulations

A previous study reported that L-PGDS contains a pocket with two binding sites, where binding site 1 includes Cys65, which functions as the catalytic residue of the enzyme.^34^ We defined this structural region as the covalent binding site of L-PGDS and ILG and used it to explore ligands targeting binding site 1. Using docking models and gel-based ABPP, we demonstrated that ILG possesses binding affinity for Cys65 of L-PGDS. Molecular modeling was performed using Yet Another Scientific Artificial Reality Application (YASARA) Structure and Maestro (Schrödinger LLC, New York, NY, USA). The 3D structure of ILG was downloaded from PubChem (https://pubchem.ncbi.nlm.nih.gov). The crystal structure of L-PGDS was obtained from the RCSB Protein Data Bank (https://www.rcsb.org/) with PDB ID 2E4J and prepared using Maestro to remove water molecules and unnecessary chains. Among the ten structural models contained in the PDB entry 2E4J, the sixth frame with the lowest RMSD was selected as the representative structure. Docking simulations were performed using YASARA Structure with the AutoDock Vina method for 25 runs. The resulting poses were automatically classified into 11 clusters using YASARA software. Cluster 3, which included Cys65 among the amino acid residues involved in binding, was selected for further analysis. In this cluster, the carbonyl moiety of ILG was oriented toward the sulfur atom of Cys65, suggesting its potential involvement in proton abstraction. The complex files (.yob) were converted into .pdb format using YASARA Structure and subsequently processed in Maestro. The Cluster3 model from YASARA was refined for docking simulations using the Protein Preparation Wizard Script in Maestro. Energy minimization of the compounds was performed using the OPLS3e force field in Maestro. A 20 × 20 × 20 Å grid box was defined by the center of mass of the ILG binding site, and the binding core was retained. The best-scoring docking pose for each hit compound was evaluated using MD simulations to assess binding stability. Binding free energies of the hit compounds were calculated using MM-GBSA (Schrödinger LLC).

### RNA isolation and reverse transcription-quantitative PCR (RT-qPCR)

RAW264.7 cells were incubated overnight in 6-well plates and treated with the indicated compounds. Total RNA was extracted from cells using ISOGEN II (Fujifilm Wako). cDNA was synthesized using the iScript Reverse Transcription Supermix for RT-qPCR (Bio-Rad) in accordance with the manufacturer’s instructions. Quantitative real-time PCR was performed with iTaq Universal SYBR Green Supermix (Bio-Rad) using the CFX Duo Real-Time PCR System (Bio-Rad). The primer sequences were designed using Primer3Plus (https://www.primer3plus.com/index.html) based on the DNA sequence obtained from NCBI (https://www.ncbi.nlm.nih.gov/) and were synthesized by Eurofins Genomics (Eurofins Scientific Group). Changes in target gene expression were calculated using the 2^-⊿⊿^ method and normalized to the housekeeping gene *Gapdh*. The relative gene, expressed as a fold-change, was calculated using the 2^−ΔΔCt^ equation. Primer sequences used for qPCR are listed in Supplemental Information Table 1.

## Data availability

All raw MS data and their peak lists are available in the jPOST repository (https://repository.jpostdb.org/) under index number JPST004464.

## Acknowledgments

This research was supported by the Japan Agency for Medical Research and Development (AMED) under Infectious Diseases Research and Infrastructure (JP25wm0325071, H.T.), AMED BINDS program for Drug Discovery and Life Science Research infrastructure project (JP25ama121029 to T.H.), the Japan Science and Technology Agency (JST) Exploratory Research for Advanced Technology (ERATO) (JPMJER2101 to H. T. and M.A.), JST FOREST program (JPMJFR230H to H.T.), JST NBDC (JPMJND2305 to H.T.), and the JSPS KAKENHI (24K02011, 24H00043, 24H00392, 24K21269, 25H01425, and 25H01426 to H.T.).

## Author Contributions

H.T. designed the study. H.S. and M.H.C. performed biological experiments. T.N. and K.O. provided insightful comments promoting this research. H.S., Y.I., M.A., K.T., and K.I. performed ABPP experiments. H.S. and T.H. performed in silico chemical-protein docking simulations. H.S. and H.T. wrote the manuscript, and all authors have thoroughly discussed this project and helped improve the manuscript.

