## Supplementary material for "Elucidation of the anti-inflammatory mechanism of isoliquiritigenin from *Glycyrrhiza uralensis* using activity-based protein profiling": Source data: Source code for Figure 1 (PCA).R

```
library(ggplot2)
library(ggprism)
library(ggnewscale)
library(ggpubr)
library(gridExtra)
library(readr)
library(plotly)
library(dplyr)
library(heatmaply)
library(tibble)
library(rstatix)
library(stringr)
library(ggbeeswarm)
library(openxlsx)
library(ggsci)
library(ggplot2)
library(RColorBrewer)

path <- "D:/2025/ABPP/results/reanalysis"
data <- read.xlsx("D:/2025/ABPP/results/reanalysis/ABPPresult_reanalysis_forR.xlsx",1)
data[is.na(data)] <- 0

xData <- data[,-c(1:3)]
metabolites <- colnames(xData)

sampleinfo <- data[,c(1:3)]


#PCA
rpca <- prcomp(x=xData,scale=TRUE)


score <- as.data.frame(rpca$x)
score$Name <- factor(sampleinfo$Name) 
order <- c("Control", "ILG 3µM", "ILG 10µM", "ILG 30µM")
score$Class <- factor(sampleinfo$Class, levels = order)
score$ILG <- ifelse(sampleinfo$Compound, "+", "-")
score$ILG <- factor(score$ILG, levels = c("-", "+"))


p12 <- ggplot(score, aes(x = PC1, y = PC2, fill = Class, shape = ILG, label = Name)) + #label = Name
  geom_point(size = 3, alpha = 0.9, color = "black") +
  scale_shape_manual(values = c(21, 22)) +
  scale_fill_manual(values = c("Control" = "#A6AEBF", "ILG 3µM" = "#3CAEA3", "ILG 10µM" = "#20639B", "ILG 30µM" = "#000B58"))+
  guides(
    fill = guide_legend(override.aes = list(shape = 21, color = "black", size = 4)),
    shape = guide_legend(override.aes = list(fill = "grey80", size = 4))
  ) +
  theme_minimal() +
  labs(
    x = paste0("PC1: ", round(summary(rpca)$importance[2,1]*100, 2), " %"),
    y = paste0("PC2: ", round(summary(rpca)$importance[2,2]*100, 2), " %")
  ) +
  theme(panel.border = element_rect(colour = "black", fill = NA, linewidth = 0.8),
        legend.text = element_text(size = 10),
        legend.position = "right", 
        panel.background = element_rect(fill="transparent", colour="transparent"),
        plot.background = element_rect(fill="transparent", colour=NA),
        aspect.ratio = 1) 
p12

svg("D:/2025/ABPP/results/reanalysis/figure/ABPP_PCA12.svg", height = 5, width =5.5, bg = "transparent")
print(p12)
dev.off()


###umap
library(umap)


data_scaled <- scale(xData)

set.seed(123)
umap_result <- umap(data_scaled)

umap_df <- as.data.frame(umap_result$layout)
colnames(umap_df) <- c("UMAP1", "UMAP2")
umap_df <- cbind(umap_df, sampleinfo)

umap_df$Name <- factor(sampleinfo$Name) 
umap_df$Class <- factor(sampleinfo$Class)
umap_df$ILG <- sampleinfo$Compound

umap <- ggplot(umap_df, aes(x = UMAP1, y = UMAP2,
                         fill = Class,
                         shape = ILG, label = Name)) +
  geom_point(size = 3, alpha = 0.9) +
  scale_shape_manual(values = c(21, 22)) +
  guides(
    fill = guide_legend(override.aes = list(shape = 21, color = "black", size = 4)),
    shape = guide_legend(override.aes = list(fill = "grey80", size = 4))
  ) +
  theme_minimal() +
  labs(title = element_blank(),
       color = "Color", shape = "Class") +
  theme(text = element_text(size = 12),
        panel.border = element_rect(colour = "black", fill = NA, linewidth = 0.8),
        plot.title = element_text(hjust = 0.5),
        aspect.ratio = 1)
umap

svg("D:/2025/ABPP/results/reanalysis/figure/ABPP_umap.svg", height = 5, width =5.5, bg = "transparent")
print(umap)
dev.off()
```
