## Supplementary material for "Elucidation of the anti-inflammatory mechanism of isoliquiritigenin from *Glycyrrhiza uralensis* using activity-based protein profiling": Source data: Source code for Figure 1 (TMT-ABPP).R

```
setwd('D:\\2025\\figure\\ABPP')
library(ggplot2)
library(openxlsx)
library(dplyr)
library(ggsignif)
library(tidyr)
library(ggnewscale)
library(patchwork)

data <- read.xlsx("D:\\2024\\ABPP\\results\\ABPP_result_forR.xlsx", 1) 
data2 <- data %>% mutate( "CR_3uM" = data$Ctrl / data$`3uM`, "CR_10uM" = data$Ctrl / data$`10uM`, "CR_30uM" = data$Ctrl / data$`30uM` )

#30uM
df_3uM <- data2 %>% .[!is.infinite(.$CR_3uM), ] %>% replace_na(list(CR_3uM = 0)) %>% .[order(.$CR_3uM,decreasing = F),] %>% mutate(rank = row_number(), class = "3uM") %>% select("rank", "Gene", "CR_3uM", "class")
#10uM
df_10uM <-  data2 %>% .[!is.infinite(.$CR_10uM), ] %>% replace_na(list(CR_10uM = 0)) %>% .[order(.$CR_10uM,decreasing = F),] %>% mutate(rank = row_number(), class = "10uM") %>% select("rank", "Gene", "CR_10uM", "class")
#3uM
df_30uM <- data2 %>% .[!is.infinite(.$CR_30uM), ] %>% replace_na(list(CR_30uM = 0)) %>% .[order(.$CR_30uM,decreasing = F),] %>% mutate(rank = row_number(), class = "30uM") %>% select("rank", "Gene", "CR_30uM", "class")

#merge　→　rbind(), cbind() (r:row, c:column)
colnames(df_3uM) <- c("rank", "Gene", "Competition_rate", "class")
colnames(df_10uM) <- c("rank", "Gene", "Competition_rate", "class")
colnames(df_30uM) <- c("rank", "Gene", "Competition_rate", "class")


plotdf <- rbind(df_3uM, df_10uM, df_30uM) %>% transform(class = factor(class, levels = c("30uM", "10uM", "3uM")))
#factor, change of the ledgend level
plotdf$class <- factor(plotdf$class, levels = c("30uM", "10uM", "3uM"))
plotdf$highlight <- ifelse(plotdf$Gene %in% c("Ptgds"), TRUE, FALSE)

###drawing the graph
g <- ggplot()+
   geom_point(data = subset(plotdf, !highlight), aes(x = rank, y = Competition_rate, color = class, shape = class))+
  annotate("rect", xmin = 2500, xmax = 2950, ymin = 2, ymax = 4, fill = "navy", alpha = 0.2, color = "navy")+
  scale_color_manual(values = c("30uM" =  "#173F5F", "10uM" =  "#20639B", "3uM" =  "#3CAEA3"))+
  scale_shape_manual(values = c("30uM" =  15, "10uM" =  17, "3uM" =  16))+
  geom_point(data = subset(plotdf, highlight), aes(x = rank, y = Competition_rate, shape = class), color = "#C5172E", size =2, show.legend = FALSE)+ 
  geom_hline(yintercept = 2, linetype = "dashed")+
  labs(x = "Peptides", y = "Competition rate")+
  scale_x_continuous(expand = c(0, 0), limits = c(0, max(plotdf$rank)*1.1), breaks=seq(0, 2500, length=6))+
  scale_y_continuous(expand = c(0, 0), limits = c(0, max(plotdf$Competition_rate)*1.05))+
  theme_classic()+
  theme(
    panel.background = element_blank(), 
    aspect.ratio = 0.8,
    legend.background = element_blank(),
    legend.title=element_blank()
  )
g


svg("ABPP_results_hl.svg", width = 4, height = 3)
print(g)
dev.off()


data3 <- subset(plotdf, Competition_rate >= 2 & Competition_rate <= 4)
data3$highlight <- ifelse(data3$Gene %in% c("Akr1b1", "Appl2", "Rhoc"), TRUE, FALSE)

g2 <- ggplot()+
  geom_point(data = subset(data3, !highlight), aes(x = rank, y = Competition_rate, color = class, shape = class))+
  scale_color_manual(values = c("3uM" =  "#3CAEA3", "10uM" =  "#20639B", "30uM" =  "#173F5F"))+
  new_scale_color()+
  geom_point(data = subset(data3, highlight), aes(x = rank, y = Competition_rate, color = Gene, shape = class), size = 2)+
  scale_color_manual(values = c("Akr1b1" = "#FE5D26", "Appl2" = "#FF9F00", "Rhoc" = "#F9CB43"))+
  labs(x = NULL, y = NULL)+
  scale_x_continuous(expand = c(0, 0), limits = c(2650, 2800))+
  scale_y_continuous(expand = c(0, 0), limits = c(2, 3.6), breaks=seq(2, 3.5, length=4))+
  theme_classic()+
  theme(
    panel.background = element_blank(), 
    aspect.ratio = 1.3, #縦/横＝1.3
    legend.background = element_blank(),
    legend.title=element_blank(),
    legend.position = "none"
  )
g2

svg("ABPP_unique.svg", width = 3, height = 3)
print(g2)
dev.off()

cmb_plot <- (g | g2) + plot_layout(heights = c(1, 1))
svg("ABPP_layout.svg", width =10, height = 5)
print(cmb_plot)
dev.off()
```
