## Supplementary material for "Elucidation of the anti-inflammatory mechanism of isoliquiritigenin from *Glycyrrhiza uralensis* using activity-based protein profiling": Source data: Source code for Figure 2 (gel-based ABPP).R

```
setwd('D:\\2025\\figure\\Gelbased-ABPP')
library(ggplot2)
library(openxlsx)
library(dplyr)
library(ggsignif)
library(viridisLite)
library(plotrix) 

###Human L-PGDS

data <- read.xlsx("D:\\2024\\Gel-based_ABPP\\results\\20241022\\241022_human_LPGDS.xlsx", 2)

order <- data$class[c(1, 4, 7, 10, 13, 16)]
data <- transform(data, class= factor(class, levels = order))

star_positions <- data.frame(
  x = c(2, 3, 4, 5, 6),
  y = c(67, 52, 28.5, 22, 28),
  label = c("**", "**", "**", "**", "***")
)

g <- ggplot(data, aes(x = class, y = ratio, fill = class))+
  stat_summary(fun = "mean", geom = "bar", width = .7, col = "black")+
  scale_fill_manual(values = c("#A6AEBF", "#B8CFCE", "#3CAEA3","#20639B", "#173F5F", "#0E2148"))+
  stat_summary(fun.max = function(x) mean(x) + std.error(x), 
               fun.min = function(x) mean(x) - std.error(x), 
               geom = "errorbar", width = .1)+
  geom_jitter(width = .1, size = 1.5)+ 
  labs(x = "", y = "Ratio (% of 0 µM)")+ 
  scale_y_continuous(expand = c(0, 0), limits = c(0, max(data$ratio)*1.1), breaks=seq(0, 100, length=5), labels = function(x) ifelse(x <= 100, x, ""))+
  theme_classic()+ 
  theme(
    axis.text.x = element_blank(), 
    axis.text.y = element_text(size = 15, color = "gray10"),
    axis.title.x = element_blank(),　
    axis.title.y = element_text(size = 15, color = "gray10"),
    legend.background = element_blank(), 
    legend.text = element_text(size = 15),
    legend.title = element_blank(), 
    plot.title = element_text(hjust = 0.5, size = 14, color = "gray10"),
    panel.background = element_rect(fill = "transparent", color = NA),
    plot.background = element_rect(fill = "transparent", color = NA),
    aspect.ratio = 1 
  )+
  geom_text(data = star_positions, aes(x = x, y = y, label = label),
            inherit.aes = FALSE, size = 5)

g

svg("gelbasedABPP_Human.svg", width = 5, height = 4, bg = "transparent")
print(g)
dev.off()


###Mouse L-PGDS
data <- read.xlsx("D:\\2024\\Gel-based_ABPP\\results\\20241022\\241022_mouse_LPGDS.xlsx", 2)

order <- data$class[c(1, 4, 7, 10, 13, 16)]
data <- transform(data, class= factor(class, levels = order))

star_positions <- data.frame(
  x = c(2, 3, 4, 5, 6),
  y = c(95, 50, 22, 20, 19.5),
  label = c("**", "**", "**", "**", "***")
)

g <- ggplot(data, aes(x = class, y = ratio, fill = class))+
  stat_summary(fun = "mean", geom = "bar", width = .7, col = "black")+
  scale_fill_manual(values = c("#A6AEBF", "#B8CFCE", "#3CAEA3","#20639B", "#173F5F", "#0E2148"))+
  stat_summary(fun.max = function(x) mean(x) + std.error(x), 
               fun.min = function(x) mean(x) - std.error(x), 
               geom = "errorbar", width = .1)+
  geom_jitter(width = .1, size = 1.5)+ 
  labs(x = "", y = "Ratio (% of 0 µM)")+ 
  scale_y_continuous(expand = c(0, 0), limits = c(0, (data$ratio)*1.1), breaks=seq(0, 100, length=5), labels = function(x) ifelse(x <= 100, x, ""))+
  theme_classic()+ 
  theme(
    axis.text.x = element_blank(), 
    axis.text.y = element_text(size = 15, color = "gray10"),
    axis.title.x = element_blank(),　
    axis.title.y = element_text(size = 15, color = "gray10"),
    legend.background = element_blank(), 
    legend.text = element_text(size = 15),
    legend.title = element_blank(), 
    plot.title = element_text(hjust = 0.5, size = 14, color = "gray10"),
    panel.background = element_rect(fill = "transparent", color = NA),
    plot.background = element_rect(fill = "transparent", color = NA),
    aspect.ratio = 1 
  )+
  geom_text(data = star_positions, aes(x = x, y = y, label = label),
            inherit.aes = FALSE, size = 5)
g

svg("gelbasedABPP_mouse.svg", width = 5, height = 4, bg = "transparent")
print(g)
dev.off()
```
