## Supplementary material for "Elucidation of the anti-inflammatory mechanism of isoliquiritigenin from *Glycyrrhiza uralensis* using activity-based protein profiling": Source data: Source code for Figure 3 (ELISA).R

```
setwd('D:\\2024\\ELISA\\20240201')
library(ggplot2)
library(openxlsx)
library(dplyr)
library(ggsignif)


data <- read.xlsx("D:\\2024\\ELISA\\20240201\\20240201_ELISA_IL6.xlsx", 5)

order <- data$class[c(1, 5, 9, 13)]
data <- transform(data, class= factor(class, levels = order))

signif_bars <- data.frame(
  x_start = c(1, 1),
  x_end = c(2, 3),
  y = c(105, 111)
)

star_positions <- data.frame(
  x = c(1.5, 2),
  y = c(106.5, 112.5),
  label = c("***", "**")
)

#######


g <- ggplot(data, aes(x = class, y = ratio, fill = class))+
  stat_summary(fun = "mean", geom = "bar", width = .7, col = "black")+
  scale_fill_manual(values = c("#F72464", "#F1EC31", "#0D92F4", "#41B3A2"))+
  stat_summary(fun.max = function(x) mean(x) + std.error(x), 
               fun.min = function(x) mean(x) - std.error(x), 
               geom = "errorbar", width = .1, position = position_dodge(width = 0.8))+
  geom_jitter(size = 1.3, width = .1)+
  labs(x = "", y = "IL-6 (% of LPS)")+ 
  scale_y_continuous(expand = c(0, 0), limits = c(0, max(data$ratio)*1.2))+
  theme_classic()+ 
  theme(
    axis.text.x = element_blank(), 
    axis.text.y = element_text(size = 12, color = "gray10"),
    axis.title.x = element_blank(),
    axis.title.y = element_text(size = 12, color = "gray10"),
    legend.key.size = unit(0.5, "cm"),
    axis.line = element_line(colour = "black"),
    legend.background = element_blank(), 
    legend.text = element_text(size = 10.5),
    legend.title = element_blank(),
    plot.title = element_text(hjust = 0.5, size = 18, color = "gray10"),
    panel.background = element_rect(fill = "transparent", color = "transparent"),
    aspect.ratio = 1 
  )+
  geom_segment(data = signif_bars, aes(x = x_start, xend = x_end, y = y, yend = y),
               inherit.aes = FALSE, size = 0.5)+
  geom_text(data = star_positions, aes(x = x, y = y, label = label),
            inherit.aes = FALSE, size = 4)

g


svg("inhibitor_ELISA_IL6_forB_v3.svg", width = 5, height = 3)
print(g)
dev.off()
```
