## Supplementary material for "Elucidation of the anti-inflammatory mechanism of isoliquiritigenin from *Glycyrrhiza uralensis* using activity-based protein profiling": Source data: Source code for Figure 3 (PGD2).R

```
setwd("D:\\2025\\figure\\PGD2")
library(ggplot2)
library(openxlsx)
library(dplyr)
library(plotrix)

data <- read.xlsx("D:\\2024\\prostaglandin_analysis\\20240605_PGD2_test_sakai.xlsx", 2)


order <- data$class[c(1, 7, 13, 19)]
data <- transform(data, class= factor(class, levels = order))

star_positions <- data.frame(
  x = c(3, 4),
  y = c(3500, 7800),
  label = c("***", "**")
)


g <- ggplot(data, aes(x = class, y = peak_height, fill = class))+
  stat_summary(fun = "mean", geom = "bar", width = .7, col = "black")+
  scale_fill_manual(values = c("#A6AEBF", "#F72464", "#173F5F", "#0D92F4"))+
  stat_summary(fun.max = function(x) mean(x) + std.error(x),
               fun.min = function(x) mean(x) - std.error(x),
               geom = 'errorbar', width = .1, position = position_dodge(width = 0.8))+
  geom_jitter( width = .1, size =　1.5)+ 
  labs(y = "ion abundance")+
  scale_y_continuous(expand = c(0, 0), limits = c(0, max(data$peak_height)*1.05))+ 
  theme_classic()+
  theme(
    axis.text.x = element_blank(),
    axis.text.y = element_text(size = 14, color = "gray10"),
    axis.title.x = element_blank(),
    axis.title.y = element_text(size = 14, color = "gray10"),
    legend.background = element_blank(),
    legend.text = element_text(size = 10),
    plot.title = element_text(hjust = 0.5, size = 18, color = "gray10"),
    panel.background = element_rect(fill = "transparent", color = NA),
    plot.background = element_rect(fill = "transparent", color = NA),
    legend.title = element_blank(),
    aspect.ratio = 1
  )+
  
  geom_text(data = star_positions, aes(x = x, y = y, label = label),
            inherit.aes = FALSE, size = 5)

g 


svg("PGD2.svg", width = 5, height = 3, bg = "transparent")
print(g)
dev.off()
```
