## Supplementary material for "Elucidation of the anti-inflammatory mechanism of isoliquiritigenin from *Glycyrrhiza uralensis* using activity-based protein profiling": Source data: Source code for Figure 3 (qPCR).R

```
library(ggplot2)
library(openxlsx)
library(dplyr)
library(ggsignif)
library(plotrix)
library(scales)

scientific_notation <- function(x) {
  x <- format(x, scientific = TRUE)
  x <- gsub("^(.*)e", "'\\1'e", x)
  x <- gsub("e", "%*%10^", x)
  x <- gsub('\\+', '', x)
  parse(text = x)
}


setwd("D:\\2025\\qPCR\\figure\\20251107")

data <- read.xlsx("D:\\2025\\qPCR\\data\\20251107\\20251107_qPCR_RAWcell.xlsx", 1)
data$Cq <- lapply(data$Cq, gsub, pattern="NaN", replacement = "") %>% as.numeric()

nrep <- data %>% distinct(Replicate) %>% .[,1]
Well <- data %>% distinct(Well) %>% .[,1]


for (i in c(1:length(Well))) {
  # i=1
  tarwell <- data %>% filter(Well == Well[[i]])
  df <- data.frame(Well = tarwell$Well[1], Target = tarwell$Target[1], class = tarwell$Biological.Set.Name[1],
                   CqMean = mean(tarwell$Cq, na.rm = T), Cq_1 = tarwell$Cq[1], Cq_2 = tarwell$Cq[2],  Cq_3 = tarwell$Cq[3])
  
  if (i==1) {
    data_mean <- df
  }
  else{
    data_mean <- rbind(data_mean, df)
  }
}


class <- data_mean %>% distinct(class) %>% .[,1]


for (i in c(1:length(class))) {
  # i=2
  Class <- class[[i]]
  tarclass <- data_mean %>% filter(class == Class)
  Cq_ref <- tarclass %>% filter(Target == "GAPDH")
  df <- tarclass %>% mutate(dCt = CqMean-mean(Cq_ref$CqMean, na.rm = T))
  
  if (i==1) {
    data_dCq <- df
  }
  else{
    data_dCq <- rbind(data_dCq, df)
  }
  
}


target <- data_dCq %>% distinct(Target) %>% .[,1]
for (i in c(1:length(target))) {
  # i=1
  # Class <- class[[i]]
  targene <- data_dCq %>% filter(Target == target[[i]])
  dCq_cali<- targene %>% filter(class == "Ctrl")
  df <- targene %>% mutate(ddCt = dCt-mean(dCq_cali$dCt, na.rm = T))
  
  if (i==1) {
    data_ddCq <- df
  }
  else{
    data_ddCq <- rbind(data_ddCq, df)
  }
}


order <- class
plotdf <- transform(data_ddCq, class = factor(class, levels = order))
plotdf <- plotdf %>% mutate("2^-ddCt" = 2^(-plotdf$ddCt)) %>% filter(Target != "GAPDH") %>% filter(class != "Ctrl")
order2 <- c("LPS","ILG", "AT56")
plotdf <- plotdf %>%  mutate(group = factor(class, levels = order2))

gene <- plotdf %>% distinct(Target)%>% .[,1]

scientific_notation <- c("1e-3", "1e-2", "1e-1")

for (i in c(1:length(gene))) {
  # i=1
  tardf <- plotdf %>% filter(Target == gene[[i]])
  tukeydf <- TukeyHSD(aov(tardf$`2^-ddCt` ~ tardf$class))
  result <- data.frame(Terget = gene[i], pair = rownames(tukeydf$`tardf$class`), tukeydf$`tardf$class`)
  
  p <- ggplot(tardf, aes(x = group, y = `2^-ddCt`, fill = class))+ 
    stat_summary(fun = "mean", geom = "bar", width = .5, col ="black")+ #alpha=0.5,col = "black", position=position_dodge2(preserve = "single")) +
    stat_summary(fun.max = function(x) mean(x) + std.error(x), 
                 fun.min = function(x) mean(x) - std.error(x), 
                 geom = 'errorbar', width = .1, position = position_dodge(width = 0.8))+
    geom_jitter(size = 1, width = .5)+ #shape = 21,col = "black", position = position_dodge(width = 0.8))+
    labs(y = expression(2^{-Delta*Delta*Ct}))+# labs(y = expression({2^-ddCt}), expression(paste("IL-1", beta)))+
    scale_fill_manual(values = c("#FF004D", "#173F5F", "#0D92F4"))+# scale_fill_manual(values = rainbow(7))+
    scale_y_continuous(expand = c(0, 0), limits = c(0, max(tardf$`2^-ddCt`)*1.2), labels = label_scientific()) + 
    theme(
      axis.text.x = element_text(size = 12, angle = 90, hjust = 1, vjust = 0.5, color= "grey10"),
      axis.text.y = element_text(size = 10, color= "grey10"),
      axis.title.y = element_text(size = 12, color= "grey10"),
      axis.title.x = element_blank(),
      legend.key.size = unit(0.5, 'cm'),
      legend.title = element_blank(),
      plot.title = element_text(hjust=0.5, size = 18, color= "grey10"),
      axis.line = element_line(colour="black"),
      panel.background = element_rect(fill = "transparent", colour = "transparent"),
      plot.background = element_rect(fill = "transparent", color = NA),
      legend.background = element_rect(fill = "transparent", color = NA),
      aspect.ratio = 1)
  
  p
  
  filepath <- paste0("D:\\2025\\qPCR\\figure\\20251107\\",gene[[i]],".svg")
  svg(filepath, width = 4, height = 3.5, bg = "transparent")
  print(p)
  dev.off()
  
  if(i ==1){
    tukey_result <- result
  }else{
    tukey_result <- rbind(tukey_result, result)
  }
  
}


write.xlsx(tukey_result, "D:\\2025\\qPCR\\data\\20251107\\tukey_result.xlsx")


###TNFa
scientific_notation_2 <- function(x) {
  x <- format(x, scientific = TRUE)
  x <- gsub("^(.*)e", "'\\1'e", x)
  x <- gsub("e", "%*%10^", x)
  x <- gsub('\\+', '', x)
  parse(text = x)
}

setwd("D:\\2025\\qPCR\\figure\\20251107")
data2_raw <- read.xlsx("D:\\2025\\qPCR\\data\\20251107\\20251107_qPCR_RAWcell.xlsx", 2)
data2_raw$Cq <- lapply(data2_raw$Cq, gsub, pattern="NaN", replacement = "") %>% as.numeric()

nrep_2 <- data2_raw %>% distinct(Replicate) %>% .[,1]
Well_2 <- data2_raw %>% distinct(Well) %>% .[,1]

for (i in seq_along(Well_2)) {
  
  tarwell_2 <- data2_raw %>% filter(Well == Well_2[[i]])
  df2 <- data.frame(
    Well = tarwell_2$Well[1],
    Target = tarwell_2$Target[1],
    class = tarwell_2$Biological.Set.Name[1],
    CqMean = mean(tarwell_2$Cq, na.rm = TRUE),
    Cq_1 = tarwell_2$Cq[1],
    Cq_2 = tarwell_2$Cq[2],
    Cq_3 = tarwell_2$Cq[3]
  )
  
  if (i == 1) data2_mean <- df2 else data2_mean <- rbind(data2_mean, df2)
}


class_2 <- data2_mean %>% distinct(class) %>% .[,1]

for (i in seq_along(class_2)) {
  
  Class2 <- class_2[[i]]
  tarclass_2 <- data2_mean %>% filter(class == Class2)
  Cq_ref_2 <- tarclass_2 %>% filter(Target == "GAPDH")
  
  df2 <- tarclass_2 %>% mutate(dCt = CqMean - mean(Cq_ref_2$CqMean, na.rm = TRUE))
  
  if (i == 1) data2_dCq <- df2 else data2_dCq <- rbind(data2_dCq, df2)
}


target_2 <- data2_dCq %>% distinct(Target) %>% .[,1]

for (i in seq_along(target_2)) {
  
  targene_2 <- data2_dCq %>% filter(Target == target_2[[i]])
  dCq_cali_2 <- targene_2 %>% filter(class == "Ctrl")
  
  df2 <- targene_2 %>% mutate(ddCt = dCt - mean(dCq_cali_2$dCt, na.rm = TRUE))
  
  if (i == 1) data2_ddCq <- df2 else data2_ddCq <- rbind(data2_ddCq, df2)
}


order_2 <- class_2
plotdf2 <- transform(data2_ddCq, class = factor(class, levels = order_2))

plotdf2 <- plotdf2 %>% 
  mutate(`2^-ddCt` = 2^(-ddCt)) %>% 
  filter(Target != "GAPDH") %>% 
  filter(class != "Ctrl")

order2_2 <- c("LPS", "ILG", "AT56")
plotdf2 <- plotdf2 %>% mutate(group = factor(class, levels = order2_2))

gene2 <- plotdf2 %>% distinct(Target) %>% .[,1]


for (i in seq_along(gene2)) {
  
  tardf2 <- plotdf2 %>% filter(Target == gene2[[i]])
  
  tukeydf2 <- TukeyHSD(aov(tardf2$`2^-ddCt` ~ tardf2$class))
  result2 <- data.frame(
    Target = gene2[i],
    pair = rownames(tukeydf2$`tardf2$class`),
    tukeydf2$`tardf2$class`
  )
  
  p2 <- ggplot(tardf2, aes(x = group, y = `2^-ddCt`, fill = class)) +
    stat_summary(fun = "mean", geom = "bar", width = .5, col = "black") +
    stat_summary(
      fun.max = function(x) mean(x) + std.error(x),
      fun.min = function(x) mean(x) - std.error(x),
      geom = 'errorbar',
      width = .1
    ) +
    geom_jitter(size = 1, width = .5) +
    labs(y = expression(2^{-Delta*Delta*Ct})) +
    scale_fill_manual(values = c("#FF004D", "#173F5F", "#0D92F4")) +
    scale_y_continuous(
      expand = c(0, 0),
      limits = c(0, max(tardf2$`2^-ddCt`)*1.2),
      labels = label_scientific()
    ) +
    theme(
      axis.text.x = element_text(size = 12, angle = 90, hjust = 1, vjust = .5),
      axis.text.y = element_text(size = 10),
      axis.title.y = element_text(size = 12),
      axis.title.x = element_blank(),
      legend.key.size = unit(0.5, 'cm'),
      legend.title = element_blank(),
      axis.line = element_line(colour="black"),
      panel.background = element_rect(fill="transparent", colour="transparent"),
      plot.background = element_rect(fill="transparent", colour=NA),
      aspect.ratio = 1
    )
  p2
  
  filepath2 <- paste0("D:\\2025\\qPCR\\figure\\20251107\\", gene2[[i]], ".svg")
  svg(filepath2, width = 4, height = 3.5, bg = "transparent")
  print(p2)
  dev.off()
  
  if (i == 1) tukey_result2 <- result2 else tukey_result2 <- rbind(tukey_result2, result2)
}

write.xlsx(tukey_result2, "D:\\2025\\qPCR\\data\\20251107\\tukey_result2.xlsx")
```
