## Supplementary material for "Elucidation of the anti-inflammatory mechanism of isoliquiritigenin from *Glycyrrhiza uralensis* using activity-based protein profiling": Source data: Source code for Figure 3 (western-blotting).R

```
setwd("D:\\2025\\figure\\WB")
library(ggplot2)
library(openxlsx)
library(dplyr)

data <- read.xlsx("D:\\2024\\WB\\result\\20231003\\excel\\20231003.xlsx", 2)


order <- data$class[c(1,5)]
data <- transform(data, class= factor(class, levels = order)) 


g <- ggplot(data, aes(x = class, y = ratio, fill = class))+
  stat_summary(fun = "mean", geom = "bar", width = .4, col = "black")+
  scale_fill_manual(values = c("#A6AEBF","#173F5F"))+#, labels = c("control", expression(paste("ILG 30", m,"M"))))+
  stat_summary(fun.max = function(x) mean(x) + sd(x), 
               fun.min = function(x) mean(x) - sd(x), 
               geom = 'errorbar', width = .1)+
  geom_jitter(width = .1, size = 1.5)+
  labs(y="Densitometric ratio of L-PGDS/GAPDH")+

  geom_text(x=1.5, y=9.5, label = "n.s.", size = 3.5, color= "grey10")+
  geom_segment(x = 1, xend = 2, y = 9, yend = 9 )+
  geom_segment(x = 2, xend = 2, y = 8.75, yend = 9) +
  geom_segment(x = 1, xend = 1, y = 8.75, yend = 9) +

  scale_y_continuous(expand = c(0, 0), limits = c(0, max(data$ratio)*1.2))+
  
  theme_classic()+
  theme(　#axis.text.x = element_blank()
    axis.text.x = element_blank(),
    axis.text.y = element_text(size = 12, color= "grey10"),
    axis.title.y = element_text(size = 10, color= "grey10" ),
    axis.title.x = element_blank(),
    legend.background = element_blank(),
    legend.text = element_text(size = 10),
    plot.title = element_text(hjust=0.5, size = 12, color= "grey10"),
    panel.background = element_rect(fill = "transparent", color = NA),
    plot.background = element_rect(fill = "transparent", color = NA),
    legend.title = element_blank(),
    aspect.ratio = 1)


g

svg("WB.svg", width = 4, height = 3, bg = "transparent")
print(g)
dev.off()
```
