## Supporting information for "Elucidation of the anti-inflammatory mechanism of isoliquiritigenin from *Glycyrrhiza uralensis* using activity-based protein profiling"

### Affiliations

### Corresponding Author

**Table S1.** Primers used for quantitative real-time PCR analysis in RAW264.7 cells.

| Gene name | Primer sequence(5'-3') |
|---|---|
| GAPDH forward | TGCCACCTTTTGACAGTGATG |
| GAPDH reverse | TGATGTGCTGCTGCGAGATT |
| L-PGDS forward | TCCGGGAGAAGAAAGCTGTA |
| L-PGDS reverse | CATAGTTGGCCTCCACCACT |
| H-PGDS forward | CACAGGAAAGGCTCTCTGCA |
| H-PGDS reverse | GTGGTGCTGCAGATATCCCA |
| IL-6 forward | AGTTGCCTTCTTGGGACTGA |
| IL-6 reverse | CAGAATTGCCATTGCACAAC |
| IL-1β forward | TGTGTCCGTCGTGGATCTGA |
| IL-1β reverse | TTGCTGTTGAAGTCGCAGGAG |
| TNFα forward | GTGGAACTGGCAGAAGAGGC |
| TNFα reverse | AGACAGAAGAGCGTGGTGGC |
| AKR1B1 forward | TGAACCAGATCGAGTGCCAC |
| AKR1B1 reverse | CATCTGGTTTGCAAGGTGCC |
| RhoC forward | CTCTCCTACCCGGACACTGA |
| RhoC reverse | CTTGGGGCTGGGAAACTCAT |
| Appl2 forward | AAGAAGATGCTGGCACCCTC |
| Appl2 reverse | TCAAGGGTTCCATCATGGCC |

**Figure S1** MS³ intensity analysis of L-PGDS in TMT-ABPP experiments.

**A.** Structure of the TMT 16-plex and an overview of its reaction mechanism with intra-protein amide groups. **B.** MS³ spectrum of L-PGDS. The isotope ion peaks from 127 to 132 are displayed in the inset.

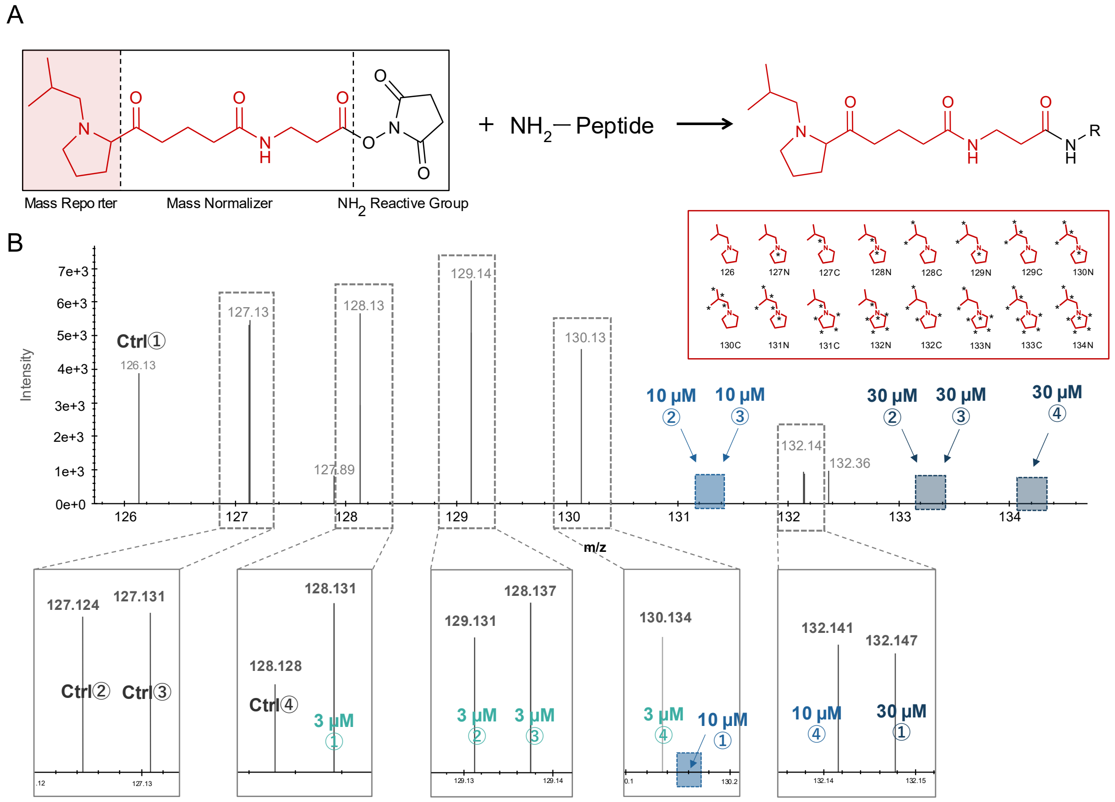

**Figure S2** Changes in gene expression levels of three candidate molecules predicted to act downstream of L-PGDS and identified by ABPP (CR > 2 ), together with the expression level of the inflammatory cytokine IL-6 measured in parallel. *P*-values were calculated relative to the LPS-treated group shown in the same graph.

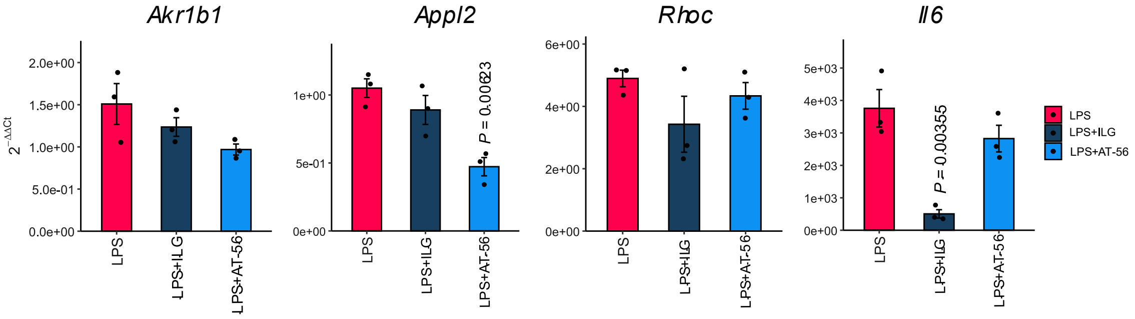
